# Temporal proteomic analysis of BK polyomavirus infection reveals virus-induced G2 arrest and highly effective evasion of innate immune sensing

**DOI:** 10.1101/601351

**Authors:** Laura G. Caller, Colin T.R. Davies, Robin Antrobus, Paul J. Lehner, Michael P. Weekes, Colin M. Crump

## Abstract

BK polyomavirus (BKPyV) is known to cause severe morbidity in renal transplant recipients and can lead to graft rejection. The simple 5.2 kilobase pair dsDNA genome expresses just seven known proteins, thus it relies heavily on host machinery to replicate. How the host proteome changes over the course of infection is key to understanding this host:virus interplay. Here for the first time quantitative temporal viromics has been used to quantify global changes in >9,000 host proteins in two types of primary human epithelial cell throughout 72 hours of BKPyV infection. These data demonstrate the importance both of cell cycle progression and pseudo-G2 arrest in effective BKPyV replication, along with a surprising lack of innate immune response throughout the whole virus replication cycle. BKPyV thus evades pathogen recognition to prevent activation of innate immune responses in a sophisticated manner.

## Introduction

BK Polyomavirus (BKPyV) is a small, non-enveloped, double stranded DNA virus that was first identified in 1971 (Gardner et al. 1971). As a ubiquitous pathogen, it establishes a life-long persistent infection in the kidneys of most humans (Viscidi et al. 2011). While infection with BKPyV is subclinical in the vast majority of individuals, it is a significant cause of morbidity in the immunosuppressed, in particular kidney and haematopoietic stem cell transplants (HSCT) recipients. Polyomavirus associated nephropathy (PVAN) affects ∼8% of kidney transplant patients, however treatment is currently limited to a reduction in immune suppression. Only a small number of anti-BKPyV drugs are available, all exhibiting significant nephrotoxicity, leading to graft decline of function and loss in ∼85% of PVAN sufferers (Huang et al. 2015). In up to 15% of HSCT patients, BKPyV leads to haemorrhagic cystitis (HC) and severely reduced rates of HSCT recovery (Arthur et al. 1986).

As with all polyomaviruses, BKPyV is structurally simple. The dsDNA genome is ∼5.2 kilobase pairs long and encodes seven proteins, three of which form the virus capsid (VP1, VP2 and VP3). The four non-structural proteins (large T antigen (LTAg), small T antigen (stAg), truncTAg and agnoprotein) have numerous functions and interact with multiple host factors. For example, LTAg binds members of the Retinoblastoma protein family, inhibiting their regulation of the G1/S phase checkpoint of the cell cycle. As a result, viral infection stimulates cell cycle progression into S phase, facilitating viral DNA genome replication (Harris et al. 1996, Stubdal et al. 1997). LTAg also binds p53, altering the regulation of both apoptosis and cell cycle progression (Lilyestrom et al. 2006). stAg modulates the phosphorylation of >300 cell cycle proteins and LTAg, through interaction with protein phosphatase 2A (Pallas et al. 1990, Scheidtmann et al. 1991). The role of agnoprotein is less well understood, although a wide range of activities have been proposed including action as a viroporin, enhancing viral DNA replication through interaction with the processivity factor proliferating cell nuclear antigen (PCNA), and enhancing egress of virions from the nucleus (Saribas et al. 2016, Panou et al. 2018).

The limited coding capacity of BKPyV necessitates co-option of multiple host factors in order to replicate and persist. Previous studies investigating how BKPyV infection modulates the host cell environment have primarily been conducted at the level of the transcriptome, which may not be reflected in the proteome. Infection in primary Renal Proximal Tubule Epithelial (RPTE) and Human Umbilical Vein Endothelial (HUV-EC) cells has been studied, either using microarray (Abend et al. 2010, Grinde et al. 2007) or RNAseq (Assetta et al. 2016, An et al. 2019). Such analyses do not provide information about virus-induced changes to cellular proteins. To date there has only been one limited analysis of changes to the host cell proteome in BKPyV infection, where stable isotope labelling with amino acids in cell culture (SILAC) was used to quantify protein changes in nuclei isolated from primary RPTE cells at 3 days post infection. In this study ∼2000 proteins were quantified, and the effect of infection on proteins outside the nucleus could not be assessed (Justice et al. 2015).

To gain a comprehensive global understanding of changes in host and viral proteins throughout the whole course of BKPyV infection, we conducted a 10-plex quantitative temporal viromic analysis (QTV) of two independent BKPyV-permissive primary human cell types, RPTE and human urothelial (HU) cells. QTV uses tandem mass tags (TMT) and MS3 mass spectrometry to quantify the relative abundance of proteins throughout the whole time course of infection (Weekes et al. 2014). These data have provided the first systematic global analysis of proteome changes caused by BKPyV infection, which has highlighted a specialised form of cell cycle arrest that is induced by this virus in primary cells. In addition, we have uncovered a complete lack of induction of innate immune responses at the protein level in BKPyV infected cells suggesting that this virus has evolved a sophisticated mechanism for evading pathogen recognition.

## Results

### Quantitative Temporal Viromic analysis of BKPyV infection

To build a global picture of changes in host and viral proteins throughout the course of BKPyV infection, we infected primary renal (RPTE) and bladder (HU) epithelial cells with BKPyV Dunlop strain. We first used 10-plex TMT and MS3 mass spectrometry to quantify changes in protein expression over three key time points of infection spanning the single-step replication cycle of this virus (0-72 hours) (Experiment 1; **Figure 1A, Figure S1A**). Cells were infected at a multiplicity of infection (MOI) of 5 infectious units per cell ensuring greater than 90% infection in both RPTE and HU cells (**Figure S1B**). In total 8985 cellular and 5/7 viral proteins were quantified in both cell lines, providing an unparalleled global view of changes in protein expression during infection in primary human epithelial cells from the kidney and bladder. Data from all proteomic experiments in this study are shown in **Table S1**. Here, the worksheet ‘‘Plots’’ is interactive, enabling generation of graphs of protein expression of any of the human and viral proteins quantified.

**Figure 1.**
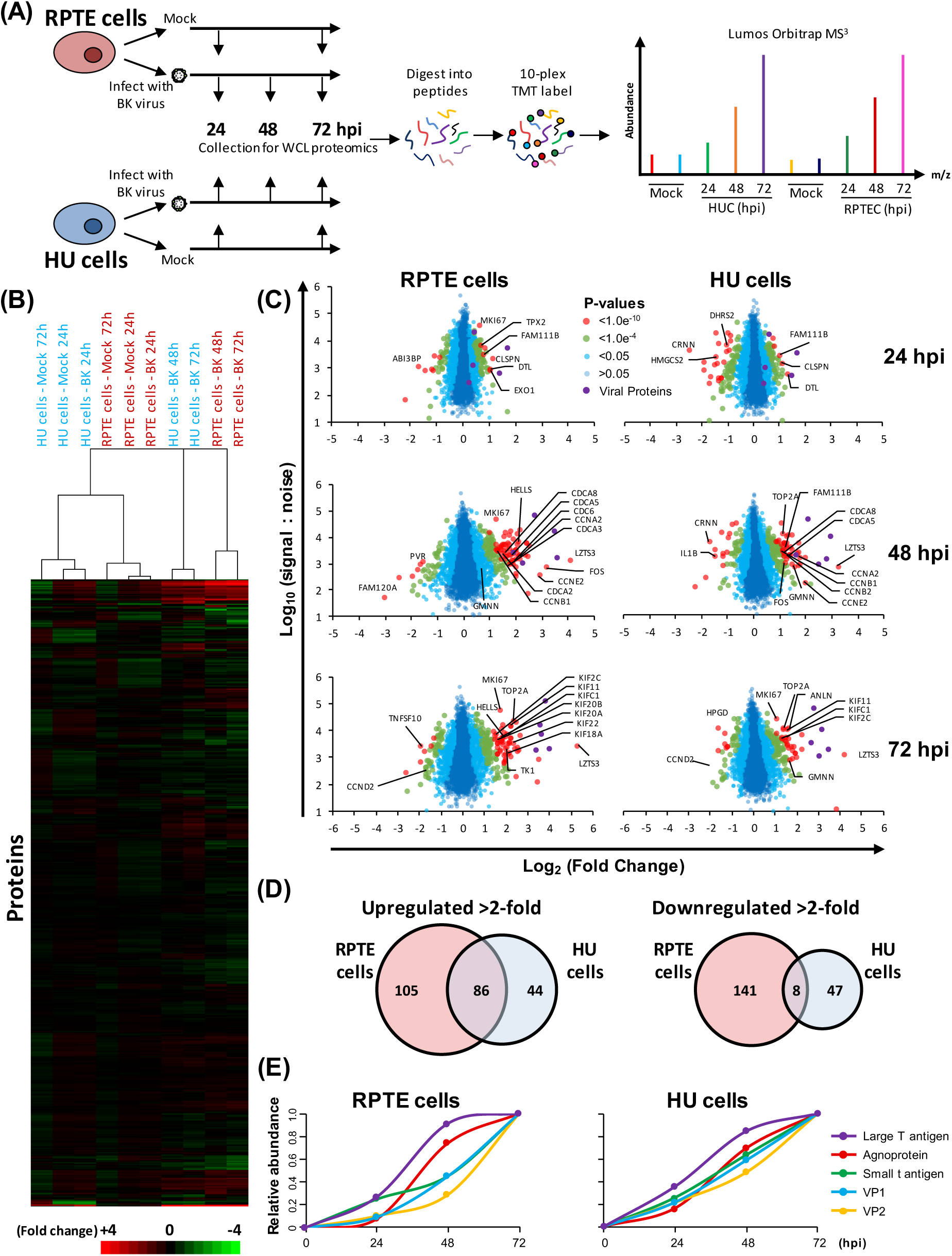
Quantitative temporal analysis of BK virus lytic infection. (A) Schematic of experimental workflow. RPTE and HU cells were infected at MOI 5 or mock infected. Whole cell lysates (WCL) were harvested at 24, 48, and 72 h (infected samples) or 24 and 72 h (mock infected samples). (B) Hierarchical cluster analysis of all quantified proteins. (C) Scatter plots of all proteins quantified at 24, 48 and 72 hpi in RPTE and HU cells. Fold change is shown in comparison to the average corresponding mock. Benjamini-Hochberg-corrected significance B was used to estimate p-values (Cox and Mann 2008). (D) Overlap of protein changes between RPTE and HU cells. (E) Temporal profiles of the 5 viral proteins identified, normalised to a maximum of one.

In uninfected cells, RPTE and HU cells exhibit differential expression of proteins, as expected from two different cell types **(Figure S2A).** In infected cells, few changes occurred by 24 h of infection, however more substantial differences were seen by 48 and 72 h (**Figure 1B-C**). In RPTE cells 191 cellular proteins increased >2-fold, while 149 proteins decreased >2-fold at any time point during BKPyV infection. In HU cells 130 proteins increased >2-fold and 55 decreased >2-fold. Many proteins showed similar changes in both cell types, although cell type-specific effects were also seen (**Figure S2**). We reasoned that those protein changes which were important for viral replication would be common to different cell types. By combining the two datasets, we found that just 86 cellular proteins, less than 1% of all proteins quantified, were upregulated >2-fold in both RPTE and HU cell lines (**Figure 1D**).

The lack of change in the host cell proteome at the earliest time point of 24 hpi suggested little or no effect of virus binding and penetration. To investigate this further a second TMT-based whole cell proteomics experiment (Experiment 2) was conducted repeating 24 and 48 hpi timepoints with an additional earlier 12 hpi timepoint, where RPTE cells were infected with UV-inactivated or unmodified BKPyV at MOI 5 (**Figures S3, S4**). Very few changes in protein abundance were observed at 12 or 24 hpi during infection with unmodified BKPyV, while at 48 hpi cellular proteins upregulated were similar to those observed in the first experiment at the same time point (**Figures S3B-D**). UV-inactivated virus induced virtually no changes at any time point, suggesting that virus replication is necessary to cause the observed changes in host protein abundance (**Figure S3B**).

### Temporal Analysis of BK Polyomavirus Protein Expression

Expression of the early BKPyV proteins, LTAg and stAg, was observed from 24 hpi, closely followed by late proteins, agnoprotein, VP1 and VP2. Profiles from HU and RPTE cells (both experiments) corresponded well (**Figures 1E, S3E**). We were unable to assign peptides to VP3 due to its 100% sequence identity with the C terminus of VP2, and the single unique peptide corresponding to the extreme N-terminus of VP3 was not quantified. Likewise, TruncTAg was not identified due to its similarity to full length LTAg: the only difference in protein sequence are the C-terminal 3 amino acids of TruncTAg, which directly follow a cluster of lysine and arginine residues and so would not be expected to be identified by our mass spectrometry analysis.

### BKPyV does not cause induction of innate immune responses in infected RPTE cells

One surprising observation from our QTV analyses was an apparent lack of an innate immune response to BKPyV infection. Of the 131 quantified proteins with annotated innate immune functions or the 69 quantified proteins with annotated antiviral functions only five were up- or downregulated 2-fold, and these changes were not consistent between the two independent cell lines or experiments (**Figure 2A and Table S2**). Despite RPTE cells being capable of mounting a response to type I interferon (**Figure 2C**), the expression of the well-established interferon stimulated genes MX1, ISG15, IFIT1, IFIT2, IFIT3, IRF3, IFI16, and BST2 remained unchanged upon BKPyV infection throughout the time course as assessed both by proteomics and immunoblot (**Figure 2B-C**). This was unexpected given that by 72 hpi large amounts of viral DNA and proteins as well as progeny virions were present within cells. This complete lack of response suggests BKPyV has evolved a highly effective immune-evasion activity, which could be due to either viral DNA and proteins not being recognised by host pathogen recognition receptors (PRRs) in these primary epithelial cells, or a potent suppression of PRR signalling pathways during BKPyV infection.

**Figure 2.**
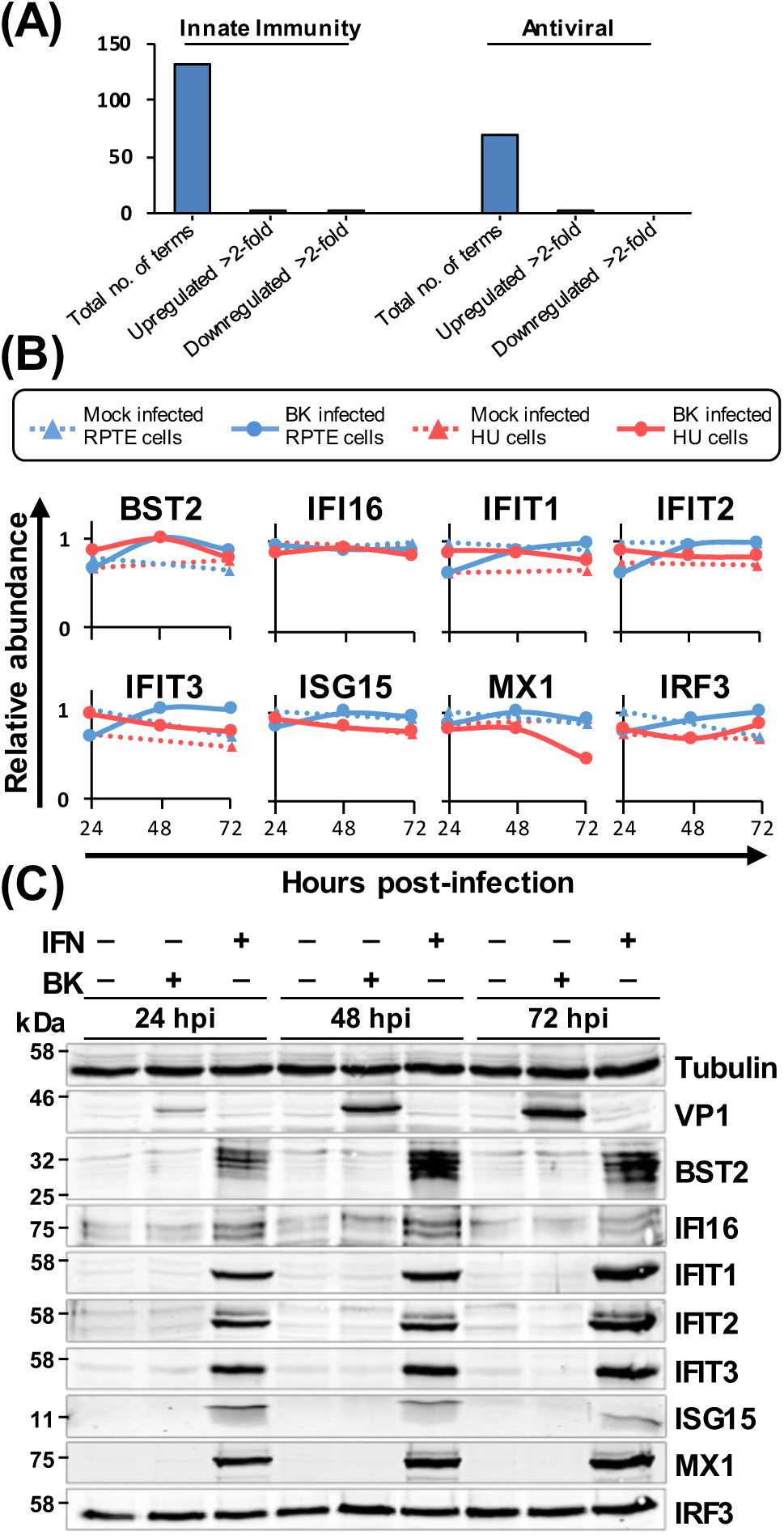
Proteins involved in the innate antiviral immune response remain unchanged during BKPyV infection. (A) Up- or down-regulation of a minority of proteins with innate antiviral function (Uniprot keywords: ‘Innate immunity’ and ‘Antiviral’). (B) Example protein profiles from (A). (C) Validation of temporal profiles shown in (B) by immunoblot (RPTE cells, MOI 3). Stimulation with IFNα2A (10^4^ U/mL) was as a positive control for innate immune response induction.

Activation of RNA and DNA sensors invariably leads to IRF3 phosphorylation and translocation into the nucleus, leading to transcription of type I and III interferons. We analysed whether RPTE cells have functional RNA and DNA sensing pathways, and whether these were activated in response to BKPyV infection. The phosphorylation and localisation of IRF3 was investigated by Western blot and immunofluorescence microscopy following BKPyV infection or treatment with Poly I:C or stimulatory DNA. Poly I:C or stimulatory DNA caused clear nuclear translocation of IRF3 in RPTE cells, with poly I:C having the greatest effect (**Figure 3A**). However, BKPyV-infected RPTE cells had no detectable change in IRF3 localisation and appeared no different to mock infected cells apart from characteristic enlarged nuclei in virus infected cells (**Figure 3A**). Furthermore, Western blot analysis showed no stimulation of IRF3 phosphorylation in BKPyV-infected RPTE cells, whereas both poly I:C and stimulatory DNA transfection caused robust IRF3 phosphorylation (**Figure 3B**). These results suggest that signal transduction pathways that would usually lead to activation of IRF3-specific kinases are not activated in infected cells either due to an inability to sense BKPyV nucleic acids or due to active inhibition by BKPyV.

**Figure 3.**
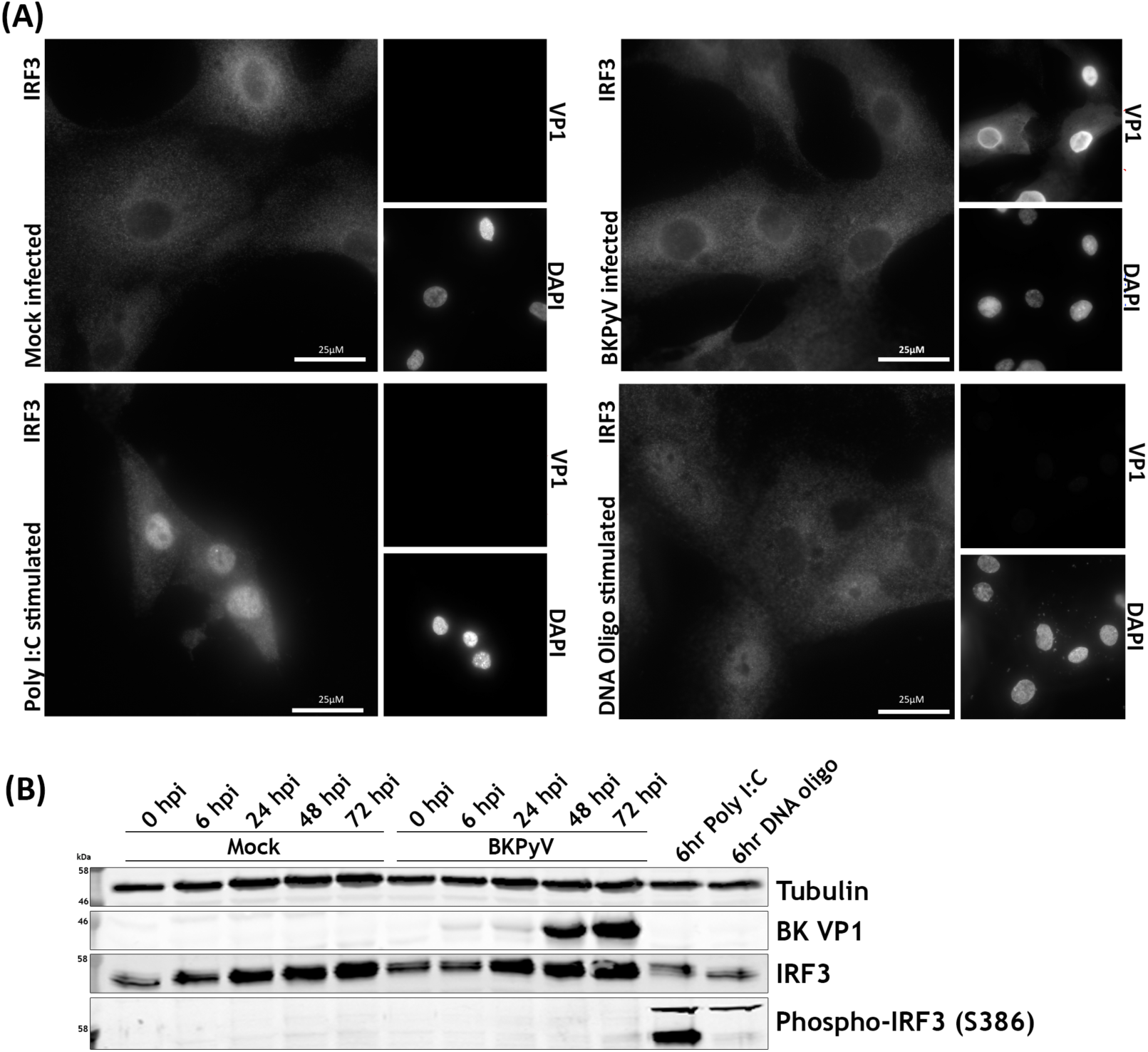
RPTE cells phosphorylate and translocate IRF3 in response to cytoplasmic RNA and DNA but fail to do so upon BKPyV infection. (A) IRF3 localisation changes upon stimulation determined by immunofluorescence microscopy. RPTE cells infected with BKPyV (MOI 0.5) or mock infected were either fixed at 48 hpi or stimulated with Poly I:C (2 µg/mL) or stimulatory DNA (2 µg/mL) and fixed at 6 h after stimulation. DAPI was used as a nuclear marker and anti-VP1 as a marker of infection. (B) Analysis of IRF3 activation by phosphorylation shown by immunoblot. RPTE cells infected with BKPyV (MOI 3) or mock infected were either fixed at 48 hpi or stimulated with Poly I:C (2 µg/mL) or stimulatory DNA (2 µg/mL) and fixed at 6 h after stimulation.

To investigate whether the lack of viral sensing is due to evasion of nucleic acid detection or active suppression of IRF3 phosphorylation, RPTE cells were mock or BKPyV-infected and subsequently stimulated with Poly I:C or stimulatory DNA at 42 hpi, prior to analysis at 48 hpi. Nuclear translocation and robust phosphorylation of IRF3 was observed in response to both RNA and DNA, irrespective of whether the cells were infected with BKPyV or mock-infected (**Figures 4A-B**). This suggests that BKPyV does not actively inhibit nucleic acid sensing pathways, IRF3 phosphorylation or IRF3 nuclear translocation. As BKPyV does not inhibit downstream activation of RNA or DNA sensing pathways this suggests BKPyV evades nucleic acid and other pathogen associated molecular pattern (PAMP) sensing pathways altogether, despite high concentrations of viral DNA, RNA and protein within these primary renal epithelial cells.

**Figure 4.**
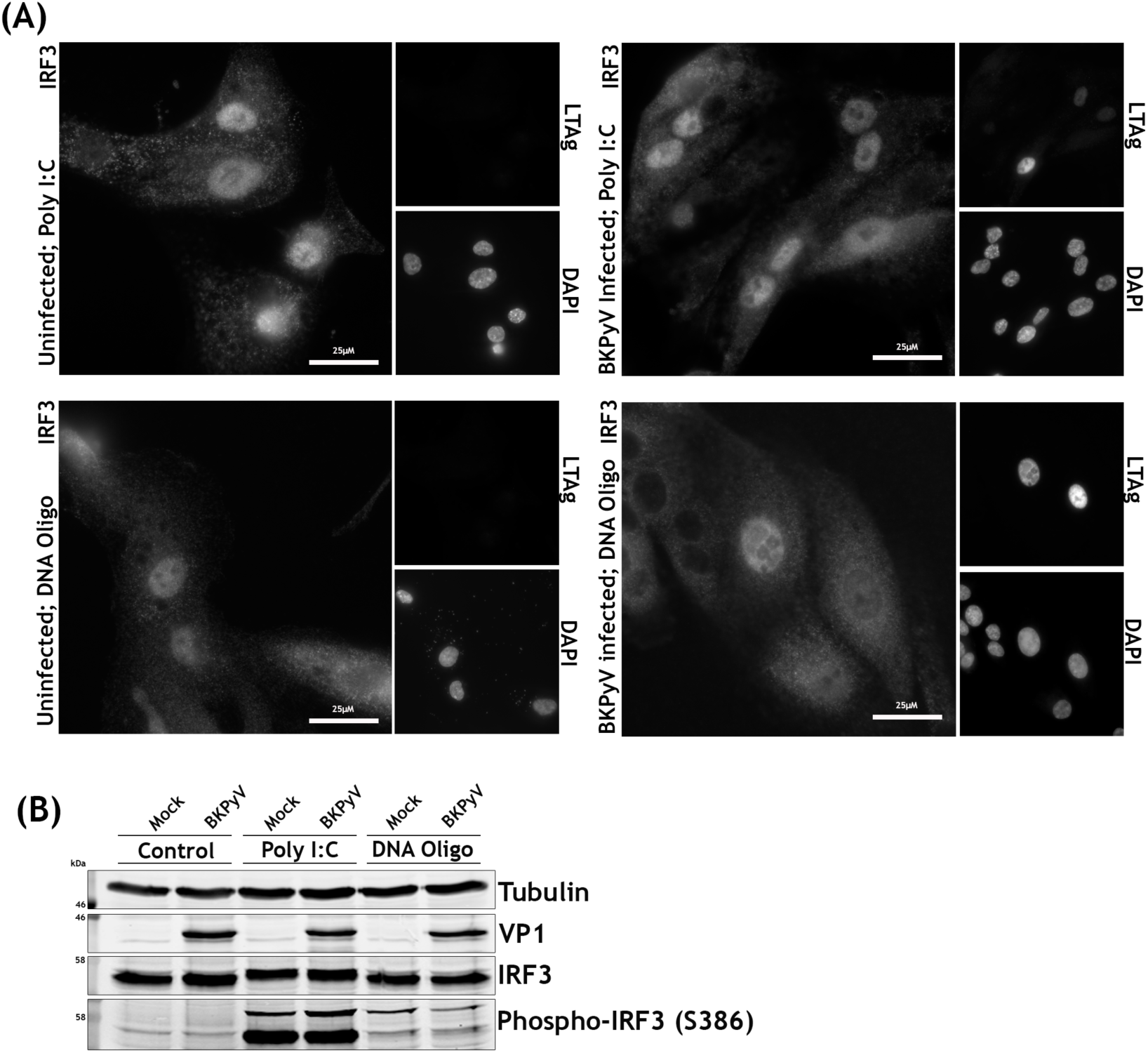
BKPyV and mock-infected RPTE cells do not differ in their responses to cytoplasmic RNA and DNA. (A) IRF3 localisation changes upon infection followed by stimulation determined by immunofluorescence. RPTE cells infected with BKPyV (MOI 0.5) or mock infected at 42 hpi were stimulated with Poly I:C (2 µg/mL) or oligomeric DNA (2 µg/mL) and fixed at 48 hpi. DAPI was used as a nuclear marker and anti-LTAg as a marker of infection. (B) Analysis of IRF3 activation by phosphorylation shown by immunoblot. RPTE cells infected with BKPyV (MOI 3) or mock infected at 42 hpi were stimulated with Poly I:C (2 µg/mL) or oligomeric DNA (2 µg/mL) and fixed at 48 hpi.

### Cell cycle associated proteins are the primary target of BKPyV

To investigate host cell functions that were modified by BKPyV, the Database for Annotation, Visualisation and Integrate Discovery (DAVID) was used to identify pathways enriched among proteins up- or downregulated during BKPyV infection (Huang et al. 2009). Amongst upregulated proteins, similar terms were enriched between HU and RPTE cells (both from experiments 1 and 2). ‘Cell cycle’ and terms related to the cell cycle dominated this analysis (**Figures 5A, S4B, Tables S3, S4**). Terms associated with the G2/M phase of the cell cycle were particularly enriched, including: chromosome, microtubule, spindle, sister chromatid cohesion and DNA damage.

**Figure 5.**
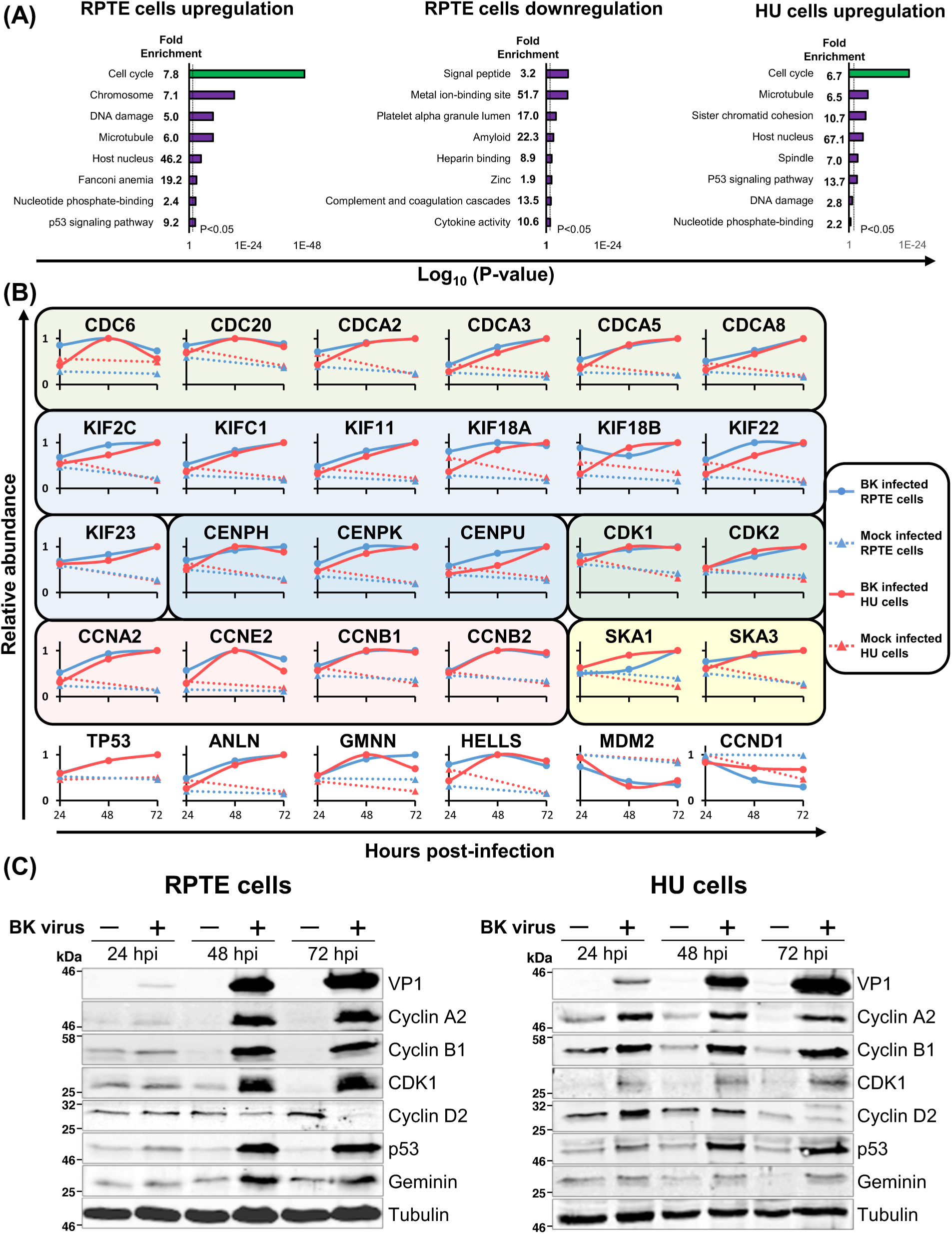
Upregulated proteins are enriched in cell cycle functions. (A) DAVID enrichment analysis of proteins upregulated or downregulated >2-fold against a background of all 8985 human proteins quantified. No significantly enriched clusters were seen in downregulated clusters in HU cells. (B) Example protein profiles from the cell cycle clusters enriched in both RPTE and HU cells. Proteins families are separated by coloured boxes. (C) Validation of temporal profiles shown in (B) by immunoblot (RPTE cells, MOI 3). Tubulin was used as a loading control and VP1 as a control for infection.

G2/M phase arrest has previously been observed in a number of different polyomavirus infections (Hornikova et al. 2017, Porras et al. 1999, Hesbacher et al. 2016, Orba et al. 2010). Cellular proteins associated with G2/M phase of the cell cycle generally increased in abundance throughout BKPyV infection including: M-phase (CDCA3), spindle formation (CDC20, CDCA2), kinetochore assembly, sister chromatid segregation and cytokinesis (KIF11, CENPK, SKA1, KIF22, ANLN), DNA repair and control of re-replication (HELLS, GMNN) and G2/M-associated cyclins and cyclin-dependent kinases (CDK1, cyclin A2 and cyclin B1) (**Figure 5B**). Proteins associated with the G1 phase, such as cyclin D2, were observed to decrease in abundance. As expected, levels of the tumour suppressor p53 were elevated during BKPyV infection; polyomavirus LTAg has a well-established function that binds, stabilises and inactivates p53 (Papadimitriou et al. 2016, Harris et al. 1996). Interestingly, MDM2, the ubiquitin ligase that normally mediates p53 degradation, was depleted during BKPyV infection (**Figure 5B**).

We confirmed these results for a number of cell cycle regulatory proteins by immunoblot throughout the time course of BKPyV infection in both RPTE and HU cells (**Figure 5B-C**). Immunofluorescence microscopy of BKPyV or mock-infected RPTE cells further confirmed the increase in cyclin B1 and CDK1 during BKPyV infection. (**Figure S5**). Furthermore, cyclin B1 remained largely cytoplasmic during BKPyV infection, suggesting that infected cells do not proceed into M phase, when cyclin B1 would normally relocalise to the nucleus (Pines and Hunter 1991).

### MDM2 and p53 levels are modulated by LTAg and cell cycle arrest

BKPyV-induced upregulation of p53 and downregulation of MDM2 was also confirmed by immunofluorescence (**Figures 6, S6**). The E3 ubiquitin ligase MDM2 is a negative regulator of both p53 and itself, leading to ubiquitinylation and degradation of p53 and MDM2 (Barak et al. 1993). In addition, p53 is a transcription factor for both itself and MDM2 (Wei et al. 2006), whose transcriptional activity is governed by the strength of extracellular and intracellular signals, such as cell cycle checkpoints, leading to the establishment of both positive and negative feedback loops (Boehme and Blattner 2009).

**Figure 6.**
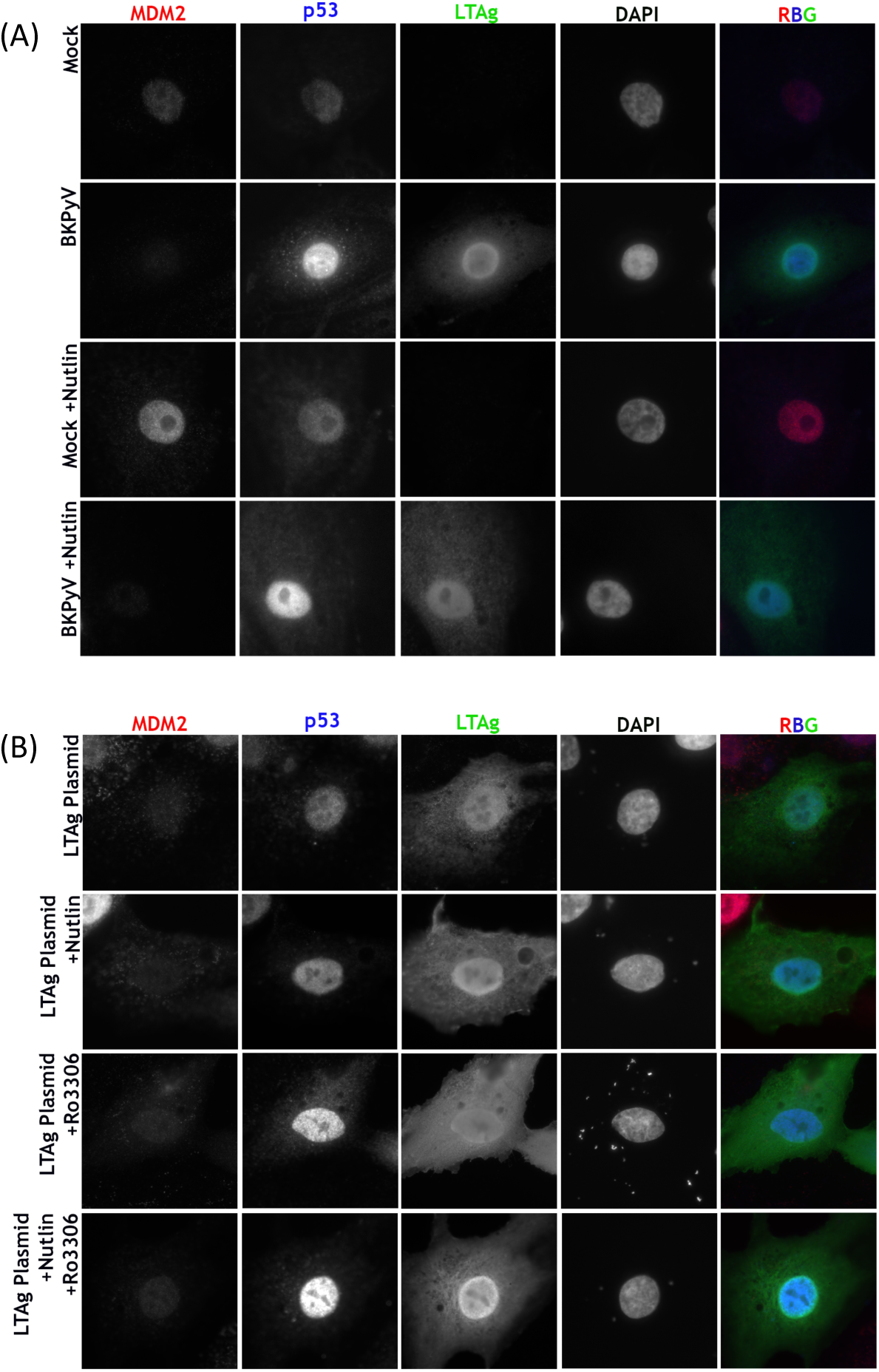
MDM2 and p53 are modulated by BKPyV via LTAg-dependent and -independent activities. (A) Expression of MDM2, p53 and LTAg in RPTE cells infected with BKPyV (MOI 1) or mock infected, then treated with 5 µM Nutlin-3 or DMSO as a control at 2 hpi, fixed at 48 hpi. DAPI was used as a nuclear marker. Select images shown, additional images provided in Figure S6. (B) Expression of MDM2, p53 and LTAg in RPTE cells transfected with BKPyV LTAg, then treated with 5 µM Nutlin-3 or DMSO as a control at 2 h and subjected to cell cycle inhibition (RO-3306 5 µM) at 24 h, fixed at 48 hpi. DAPI was used as a nuclear marker. Select images shown, additional images provided in Figure S6.

Polyomavirus LTAgs are well established to efficiently bind, stabilise and inhibit p53 (Sheppard et al. 1999), leading to increased p53 levels in BKPyV infected cells. Our data now demonstrates for the first time that this is accompanied by a decrease in MDM2 levels. Interplay between BKPyV infection, MDM2 and p53 was investigated using the MDM2 inhibitor Nutlin-3. Nutlin-3 occupies the p53 binding pocket on MDM2 obstructing their interaction and leading to reduced p53 ubiquitinylation. In addition, Nutlin-3 leads to increased transcription of MDM2 due to the release of active p53 (Vassilev et al. 2004). Mock or BKPyV infected RPTE cells were treated with Nutlin-3 at 2 hpi, or DMSO as a control, and fixed at 48 hpi. Cells were immunostained for expression of MDM2, p53, and LTAg (infection marker) (**Figure 6A**). Low endogenous levels of both MDM2 and p53 were observed in the nuclei of untreated mock infected cells. BKPyV infection lead to a reduction in MDM2, while p53 substantially increased, correlating with the changes observed in the proteomics data. Mock infected cells treated with Nutin-3 showed increased levels of MDM2, accompanied by a slight increase in p53 levels, in accordance with published effects of Nutlin-3 (Vassilev et al. 2004). Interestingly, MDM2 levels did not increase in BKPyV infected cells treated with Nutlin-3 and in fact MDM2 levels were observed to reduce, whilst p53 levels were once again substantially increased. This suggests that during infection MDM2 remains able to self-ubiquitinylate, leading to its degradation in the presence of Nutlin-3, however transcription of MDM2 by p53 is apparently inhibited, likely due to p53 sequestration by LTAg.

RPTE cells were transfected with LTAg to investigate whether LTAg expression alone was sufficient to cause the observed MDM2 decrease and p53 increase. At 2 h cells were treated with Nutlin-3 or DMSO and then in addition some samples were treated at 24 h with a CDK1-specific inhibitor, RO-3306, to simulate BKPyV induced cell cycle arrest. Cells were fixed at 48 h and immunostained for MDM2, p53, and LTAg (**Figure 6B**). Transfection of LTAg alone was sufficient to reduce MDM2 levels, however p53 levels were increased only slightly suggesting other effects of BKPyV infection in addition to LTAg expression modulate p53 and MDM2 levels. Nutlin-3 treatment did not alter the effects of LTAg on MDM2 or p53 levels. Treatment of LTAg transfected cells with the CDK1 inhibitor RO-3306 lead to a marked increase in p53 expression, while MDM2 levels were once again decreased. Combined Nutlin-3 and RO-3306 treatment further enhanced the increase of p53 in LTAg expressing cells. Taken together, these data suggest LTAg binding to p53 displaces MDM2 leading to p53 stabilisation and MDM2 degradation, but LTAg binding also prevents p53-dependent expression of MDM2 and p53. Furthermore, virus infection or G2/M arrest stimulates p53 expression, possibly via a DNA damage-type response.

### BKPyV induced G2/M phase arrest is prevented by inhibition of CDK1 & CDK2, but not by inhibition of CDK1 alone or CDK4 & CDK6

Given the striking dysregulation of cell cycle-related proteins during BKPyV replication, we postulated that BK-induced pseudo-G2 phase may serve a number of roles in BKPyV replication. We therefore investigated the effect of BKPyV infection on the host cell cycle status in the presence or absence of various CDK inhibitors. Polyomavirus replication is heavily reliant on the host DNA synthesis machinery and it has previously been shown that either BKPyV infection or JCPyV LTAg expression alone can cause cells to arrest in the G2/M phase of the cell cycle (Jiang et al. 2012, Verhalen et al. 2015, Orba et al. 2010). However, the impact of CDK inhibitors on BKPyV-induced arrest has not been fully investigated. RPTE cells were infected with BKPyV, treated with CDK inhibitors after 24 hpi to allow sufficient time for virus entry and initiation of early gene expression, then subsequently harvested at 48 hpi and analysed by flow cytometry to compare cell cycle profiles. PD0332991 was used to inhibit CDK4 and 6, which are active in G1 phase, Roscovitine was used to inhibit CDK1 and 2, which are active throughout S, G2, and M phase, and RO-3306 was used to inhibit CDK1, which is active in G2 and M phase. In mock-infected RPTE cells, all three inhibitors produced the expected effects: increased G1 for PD0332991; increase G2/M for RO-3306; no change for Roscovitine (**Figure 7**). Infection of RPTE cells with BKPyV in the absence of any inhibitor increased the proportion of cells in G2/M and S phases from ∼28% (mock) to ∼42% (BKPyV), consistent with previous published data (Jiang et al. 2012). BKPyV-infected cells that were treated with PD0332991 showed a slight increase in the proportion of cells in G1 and S phases compared to control infected cells, although these samples still showed substantially more cells in G2/M than uninfected cells. This suggests that inhibition of CDK4 and 6 does not prevent BKPyV driving infected cells through the G1/S checkpoint, due to Rb inactivation by LTAg, or arresting cells in G2/M. In contrast, treatment of infected cells with Roscovitine, which inhibits both CDK1 and 2, appear to severely restrict BKPyV-stimulated S phase entry and G2/M arrest, as the cell cycle status was similar to that of mock-infected cells with >72% of cells in G1, ∼8% in S phase, and ∼19% in G2/M phase (**Figure 7A**). Infected cells treated with RO-3306 (CDK1 inhibitor) showed a similar cell cycle profile to that of control-treated infected cells, (**Figure 7A**). This suggests that inhibition of CDK1 alone does not affect the ability of BKPyV to induce progression through S phase and subsequent G2/M arrest, and further that BKPyV infection and CDK1 inhibition have similar effects on the cell cycle, namely inducing a G2/M arrest.

**Figure 7.**
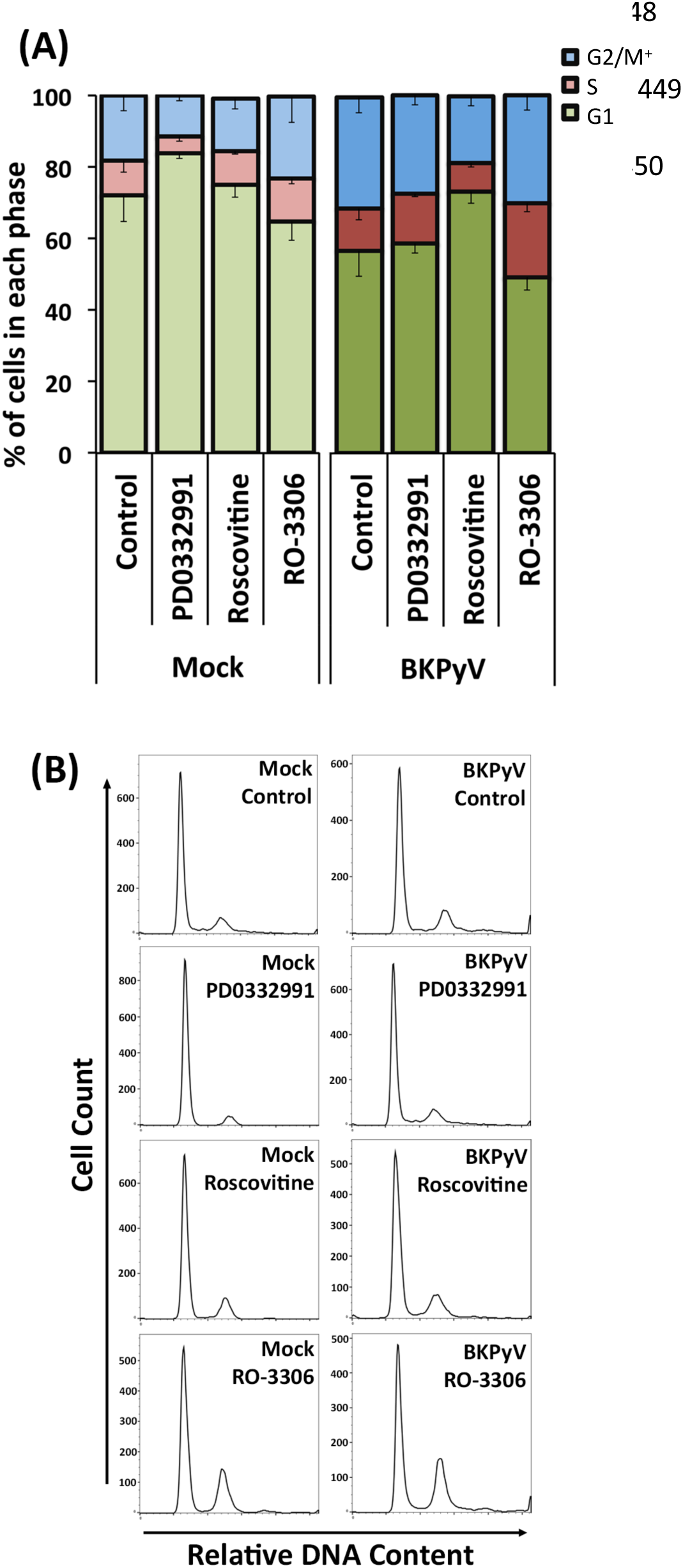
Cell cycle inhibitors have variable effects on BKPyV induced G2/M phase cell cycle arrest. (A) The cell cycle status of RPTE cells was determined in a number of different experimental conditions. RPTE cells were infected with BKPyV (MOI 3) or mock infected, then subjected to CDK4/6 inhibition (PD0332991 1 µM), CDK1/2 inhibition (Roscovitine 20 µM) or CDK1 inhibition (RO-3306 5 µM) at 24 hpi, and subsequently collected for analysis at 48 hpi. Collected cells were stained with propidium iodide (PI) and analysed by flow cytometry (n=3). Error bars represent standard deviation. (B) Histograms of PI stain for each experimental condition of a single experiment shown.

### Inhibition of CDK1 and CDK2 or CDK1 alone reduces BKPyV replication

The ability of BKPyV to induce a pseudo-G2 arrest in the presence of CDK1 or CDK4/6 inhibition suggested that virus replication should be unaffected in such conditions, while inhibition of CDK1/2 should perturbed viral replication due to inhibition of S phase progression. To investigate if this was the case we next analysed the effect of CDK inhibitors on viral genome synthesis in BKPyV infected RPTE cells. Infected cells were again treated with each inhibitor at 24 hpi and harvested at 48 hpi. Viral and host cell DNA was quantified using qPCR to determine viral DNA copy numbers per cell and were normalised to uninhibited controls (arbitrarily set to 1). Inhibition of CDK4/6 had no significant effect on viral genome synthesis, while inhibition of CDK1 and 2 by Roscovitine showed a 7.4-fold reduction in the synthesis of BKPyV genome, likely due to the restriction of cells from entering and progressing through S phase (**Figure 8A**). Surprisingly inhibition of CDK1 alone by RO-3306 also caused a significant, although more modest, 2.3-fold reduction in BKPyV genome synthesis. This suggests that, despite this inhibitor having little effect on BKPyV-driven cell cycle progression, CDK1 activity is important for efficient viral genome synthesis.

**Figure 8.**
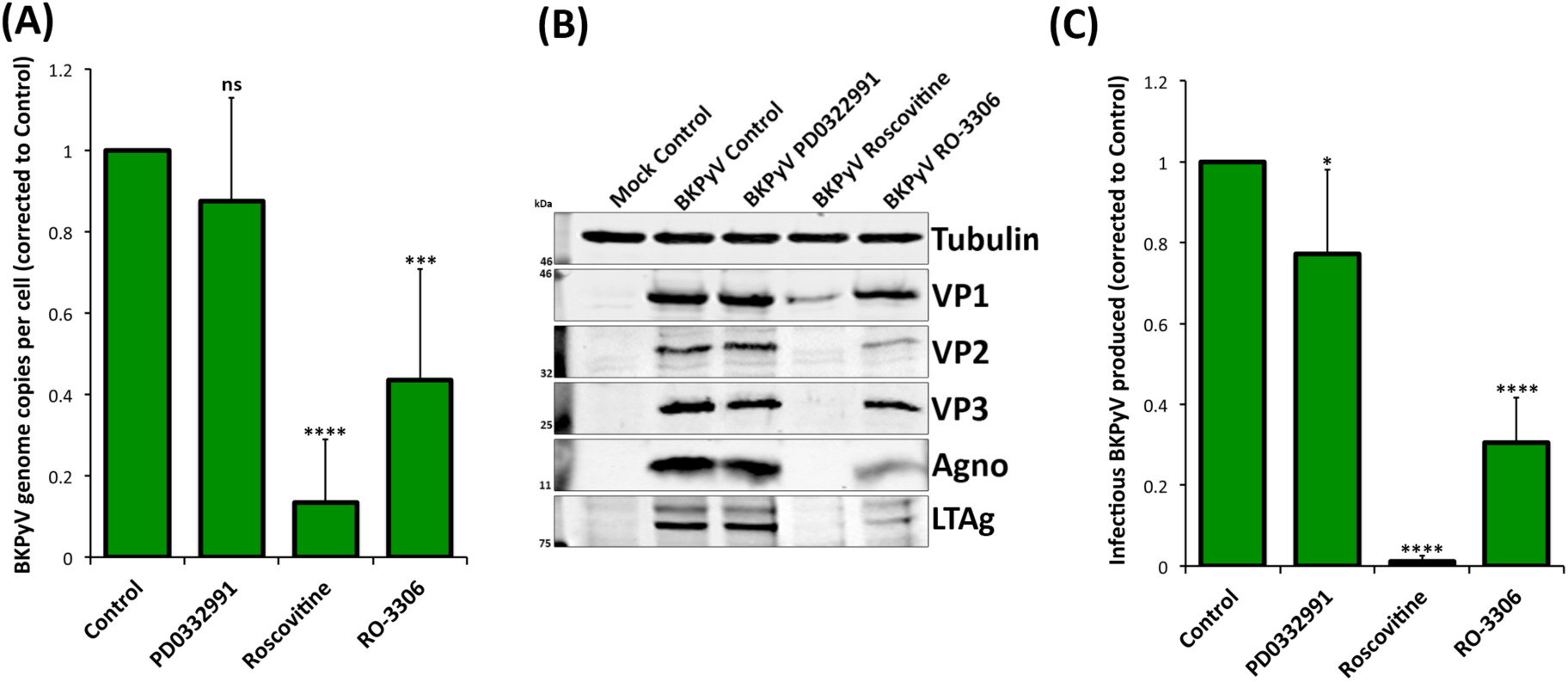
CDK1 and 2 inhibitors impede BKPyV replication in RPTE cells. RPTE cells infected with BKPyV (MOI=3) were subjected to CDK4/6 inhibition (1 µM PD0332991), CDK1/2 inhibition (20 µM Roscovitine) or CDK1 inhibition (5 µM RO-3306) from 24 hpi and harvested for analysis at 48 hpi. (A) qPCR to determine viral DNA copy numbers per cell. DNA was extracted from each condition, BKPyV genome copy number was determined, normalised to host gene (TNFα) copy number and compared to the uninhibited control, which was arbitrarily set to 1 (n=6). (B) Expression of viral proteins VP1, VP2, VP3, Agno, and LTAg was determined by immunoblot. Tubulin was used as a loading control. (C) Infectious BKPyV produced in each experimental condition was determined by fluorescent focus unit (FFU) assay and normalised to uninhibited control (arbitrarily set to 1) (n=7). Error bars represent standard deviation. *p < 0.05; ***p < 0.001; ****p < 0.0001; ns = not significant; one sample t-test experimental conditions versus control.

The effects of these CDK inhibitors on viral protein synthesis was similarly investigated (**Figure 8B**). Inhibition of CDK4 and 6 had no observable effect on viral protein synthesis, while inhibition of CDK1 and 2 by Roscovitine substantially reduced viral protein expression levels. Inhibition of CDK1 alone by RO-3306 showed only small reductions in viral protein levels (**Figure 8B**).

Analysis of infectious virus production in the presence or absence of these CDK inhibitors also demonstrated a similar trend. Inhibition of CDK4 and 6 caused only a slight reduction of infectious titres, whereas inhibition of CDK1 and 2 by Roscovitine resulted in a significant 80-fold reduction of virus production, unsurprisingly given the dramatic inhibition of viral DNA and protein synthesis (**Figure 8C**). Inhibition of CDK1 alone by RO-3306 caused a significant reduction of infectious virus titre by >3-fold. These data further suggest that CDK1 activity is important for the efficient production of infectious viruses.

## Discussion

By employing the power and sensitivity of TMT-based MS3 mass spectrometry technology, we have been able to uncover BKPyV-induced changes to protein abundance and an unparalleled global view of protein expression within these human cell types. Importantly, these studies were conducted in primary human cells from epithelial tissue representing the natural sites of replication *in vivo*. Therefore, this work also provides a comprehensive proteomic resource for future studies on human renourinary epithelial biology.

One of the most surprising findings of this study was just how few of the ∼9000 cellular proteins that were quantified changed in abundance in response to BKPyV infection. In fact, just 235 were found to be upregulated and 196 downregulated >2-fold or more across either cell line at any time point, which corresponds to <5% of the total proteome. Previous studies that applied a similar TMT-based approach to infection with human cytomegalovirus (HCMV), another dsDNA virus, revealed that 56% of cellular proteins changed in abundance more than 2-fold during the course of infection (Weekes et al. 2014). This suggests that BKPyV, and presumably other polyomaviruses, are so highly adapted to their host that they only need to induce subtle changes to host gene expression to very effectively reprogram cells into virus-producing factories. This also suggests polyomaviruses can very effectively evade detection by host pathogen recognition receptors despite producing high concentrations of foreign (viral) nucleic acid and proteins seen in high multiplicity infections such as these.

For host proteins induced by BKPyV infection, we identified substantial overlap between the two primary cell lines, with many of the same or highly related functional clusters identified by DAVID analysis. This includes clusters such as ‘DNA damage’ and the ‘Fanconi Anaemia’ pathway, which have been previously described as important during polyomavirus replication to ensure viral genome replication maintains high fidelity (Jiang et al. 2012). Interestingly, the majority of the functional clusters identified as upregulated in BKPyV infection are related to cell cycle activity and regulation, in particular activities associated with G2 and M phases. In fact, BKPyV infection appears to have a similar ‘G2/M arrest’ effect on cell cycle status as the CDK1-specific inhibitor RO-3306, a drug commonly used to arrest cells in G2. The fact that infected cells do not progress into authentic mitosis is supported by the observation that cyclin B1 remains predominantly cytoplasmic despite higher protein levels in infected cells, and supports previously published data indicating G2/M phase arrest is driven by polyomavirus infection (Harris et al. 1996, Lilyestrom et al. 2006, Stubdal et al. 1997, Rundell and Parakati 2001, Orba et al. 2010). Importantly our data provides a much greater understanding of host protein profiles that are associated with polyomavirus-induced G2 arrest, and it would be interesting to compare these observations to the effects of RO-3306 or other specific CDK1 inhibitors on cellular protein expression profiles.

Our data also indicate a specific perturbation of the p53-MDM2 axis by BKPyV infection, where MDM2 is destroyed and p53 is substantially increased but kept inactive by LTAg binding. However, these changes require more than just LTAg expression, and are also driven by additional effects of BKPyV infection related to G2/M arrest and potentially DNA damage responses. Our findings suggest the following model: low MDM2 and p53 levels are maintained in uninfected cells due to their poly-ubiquitinylation by MDM2 and subsequent proteosomal degradation (**Figure 9A**). Inhibition of MDM2-p53 interaction by Nutlin-3 releases p53 which then stimulates MDM2 expression (**Figure 9B**). Interaction of LTAg with p53 displaces MDM2, thereby causing MDM2 to be destroyed by the proteasome, and so protects p53 from degradation but inhibits p53 transcriptional activity prevent induction of de novo MDM2 expression (**Figure 9C**). Therefore, the expression of just LTAg results in decreased MDM2 levels but only a modest increase in p53. During active BKPyV infection, p53 expression is induced by some other effect(s) of virus replication, and these additional copies of p53 are also bound and inactivated by LTAg (**Figure 9D**). The virus induced stimulation of p53 expression is predicted to be via a DNA damage response pathway, which can be mimicked by inducing a G2 arrest through inhibition of CDK1.

**Figure 9.**
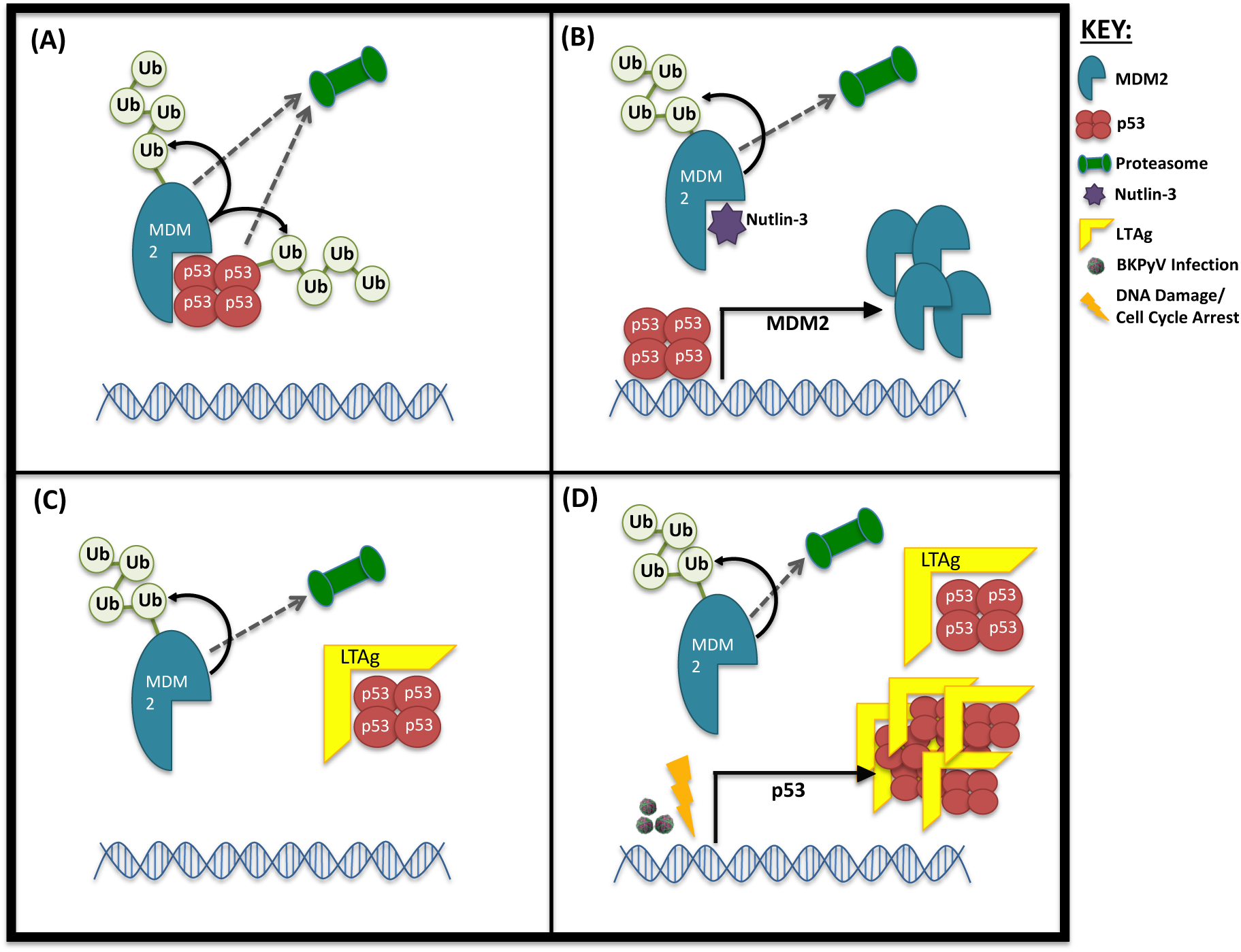
Proposed mechanism of interplay between MDM2 and p53 levels in the presence of LTAg. (A) Untreated cells. (B) Uninfected, untransfected cells inhibited with Nutlin-3. (C) Cells expressing LTAg in the absence of infection. (D) Cells BKPyV infected, or cells expressing LTAg combined with cell cycle arrest/DNA damage.

Polyomaviruses have a well-established capacity to drive cells into S-phase by overriding the G1/S checkpoint via the activity of LTAg. It is therefore unsurprising that inhibition of CDK4 and 6 has little-to-no effect on the ability of BKPyV to drive cell cycle progression or to replicate. CDK4 and 6, in complex with cyclin D, are normally responsible for phosphorylation of Rb and release of E2F proteins allowing passage through the G1/S checkpoint (Chellappan et al. 1991). This is bypassed through the binding of LTAg to Rb family proteins, releasing E2F proteins enabling S-phase entry that is unconstrained by upstream factors (Harris et al. 1996, Stubdal et al. 1997).

In stark contrast Roscovitine, a potent inhibitor of CDK1 and 2, by caused a global cell cycle arrest, irrespective of BKPyV infection, and drastically reduced BKPyV replication. Similar effects of Roscovitine on polyomavirus replication have been previously been attributed to inhibition of CDK1 activity alone (Orba et al. 2008). However, our data suggests the effect of Roscovitine are more likely due to inhibition of CDK2 activity, or the combination of inhibiting both CDK1 and 2. Inhibition of both these cyclin dependent kinases causes a rather global block to cell cycle progression; CDK2 is active in both late G1 and S phase, while CDK1 is active in G2 and M phase (Obaya and Sedivy 2002). CDK2 activity is required immediately after G1 checkpoint clearance and beyond, and so the primary cause of BKPyV inhibition by Roscovitine could be due to a failure to activate S-phase proteins required for viral genome synthesis and consequent protein expression.

Somewhat more intriguing is the effect of CDK1-specific inhibition on BKPyV infection; RO-3306 caused significant reductions in viral DNA synthesis and infectious virus assembly. This was surprising because CDK1 activity is normally important for the transition through G2 and into M-phase, and so inhibition of CDK1 would not be expected to inhibit progression through S-phase and thus viral DNA replication. Whether CDK1 activity is required directly or indirectly to enhance DNA synthesis or other S-phase activities required for BKPyV genome replication, or the process of virion assembly, remains to be determined.

Moreover, our data have demonstrated that BKPyV infection of renourinary epithelial cells does not appear to cause the induction of antiviral responses in agreement with published transcriptome data of RPTE cells (An et al. 2019). Both RPTE and HU cells express the appropriate receptors, signalling pathways and transcription factors associated with sensing and responding to DNA viruses, such as cGAS (MB21D1), IFI16, STING (TMEM173), NFκB, and IRF3, which were readily detected in our mass spectrometry analysis (**Table S1**). RPTE cells are quite capable of responding to foreign intracellular DNA or RNA, leading to phosphorylation and nuclear translocation of IRF3, and RPTE cells also robustly express antiviral genes in response to type-1 interferon. However, we could not detect activation of these pathways even by three days after BKPyV infection: IRF3 remains unphosphorylated and cytoplasmic and no ISGs were induced. Furthermore, we have also shown that active BKPyV infection within the same cell does not prevent the phosphorylation and nuclear translocation of IRF3 in response to either cytoplasmic RNA or DNA. This suggests that BKPyV is not actively suppressing such antiviral responses, but rather prevents its own detection by pathogen recognition receptors. This evasion of detection may be due to a combination of having a small circular double stranded DNA genome that is associated with histones, thus appearing similar to open chromatin, and tightly regulated entry and egress mechanisms to prevent exposure of viral DNA in the cytoplasm. Whether an inability to sense and respond to BKPyV infection is partly due to the nature of epithelial cells in the renourinary systems and whether this contributes to the natural tropism of BKPyV for these tissue types will be interesting questions for future study.

In summary, we have generated extensive data sets on the protein expression profiles of primary epithelial cells of the kidney and bladder using advanced multiplexed proteomics and provided a detailed understanding of how infection by BKPyV modifies the protein expression profiles in these cells. This has uncovered a highly specific cell cycle arrest induced by virus infection and revealed the importance of this arrest for BKPyV replication, as well as uncovering details of p53-MDM2 antagonism by this virus. Furthermore, our findings suggest a surprising ability of BKPyV to evade detection and activation of innate immune responses in cells that are natural sites of lytic virus infection *in vivo*.

## Materials and Methods

### Cell lines, virus and primary antibodies

HU cells were grown in Urothelial Cell Medium enriched with Urothelial Cell Growth Supplement and penicillin/streptomycin solution (Caltag Medsystems). HU cells were used at passage 4-6 for all experiments. RPTE cells were grown in renal epithelial basal media enriched with REGM bullet kit (Lonza). RPTE cells were used at passage 6-7 for all experiments.

BKPyV (Dunlop strain) inserted into pGEM7Zf(+) vector (kindly provided by M. Imperiale, University of Michigan) was digested with BamHI, purified and re-ligated. Resultant BKPyV-Dunlop genome was transfected into a T150 flask of RPTE cells, one week later the flask was split into three T150 flasks of RPTE cells. After a period of up to four weeks virus was harvested by freeze thawing cells three times. Virus purification by sucrose cushion, followed by caesium chloride gradient and dialysis provided purified BKPyV stocks as described previously (Jiang et al. 2009). Concentration and purity was assessed by FFU assay and coomassie gel stain respectively.

The primary antibodies used in this study were PAb597 against SV40 VP1 (kindly provided by W. Atwood, Brown University), P5G6 against BKPyV VP1 (kindly provided by D. Galloway, Fred Hutchinson Cancer Research Center), ab6160 against Tubulin (Abcam), ab32386 against Cyclin A2 (Abcam), ab32053 against Cyclin B1 (Abcam), MA5-11472 against CDK1 (Thermo Scientific), ab207604 against Cyclin D2, ab1101 against p53 (Abcam), GTX116125 against Geminin (GeneTex), ab16895 against MDM2 (Abcam), 37849 against MX1 (Cell Signalling Technologies), 2758 against ISG15 (Cell Signalling Technologies), PA3-848 against IFIT1 (ThermoFisher), 12604-1-AP against IFIT2 (ProteinTech), SAB1410691 against IFIT3 (Sigma Aldrich), 11904 against IRF3 (Cell Signalling Technologies), ab76493 against IRF3 (phospho S386) (Abcam), ac-8023 against IFI16 (Santa Cruz), 11721 against BST2 (NIH AIDS Reagent Programme), ab53983 against SV40 VP2 + VP3 (Abcam), ab16879 against SV40 LTag (Abcam), and against BKPyV agnoprotein (rabbit polyclonal antibody generated against agnoprotein specific peptide).

### Cell infections and harvesting virus

For viral infections RPTE or HU cells were infected with BKPyV at either MOI=5 for TMT and validation experiments, or MOI=3 or 0.5 for all other experiments, diluted in appropriate medium. At 1 hpi media was removed, cells washed twice with PBS and fresh medium was added. For TMT analysis, cells were harvested in TMT lysis buffer (6M Guanidine HCl, 50mM HEPES pH 8.5), vortexed extensively and incubated at room temperature for 10 min. Lysates were then sonicated at 25 W for 30 s, followed by centrifugation at 21,000 g for 10 min, after which supernatant was transferred to a fresh tube. Centrifugation was repeated and supernatants snap-frozen in liquid nitrogen for further processing. For immunoblot, cells were harvested by centrifugation at 6,000 g after two PBS washes.

### Transfection

RPTE cells were transfected with pcDNA3-LTAg plasmid using TransIT-LT1 Transfection Reagent (Mirus) in Opti-MEM media according to the manufacturers protocol.

### Inhibitors

For p53:MDM2 interaction inhibition experiments cells were treated at 2 hpi with Nutlin-3. Nutlin-3 (Sigma) was made up to 20mM in DMSO and used at 5µM. For cell cycle inhibition experiments cells were treated with inhibitors at 24 hpi. PD0332991 (Sigma) was made up to 5mM in dH_2_O and used at 1µM, Roscovitine (Sigma) was made up to 20mM in DMSO and used at 20µM and RO-3306 (Sigma) was made up to 20mM in DMSO and used at 5µM. Controls were subjected to treatment with an equivalent amount of DMSO at the greatest volume of any inhibitor used. Cells were harvested in 1mL media at 48 hpi and either pelleted by centrifugation at 6,000 g for use in western blot or qPCR, or frozen for assay by FFU. For analysis by flow cytometry cells were detached from wells by trypsin/EDTA treatment, centrifuged at 6,000 g, washed in PBS and fixed in 70% ice-cold ethanol.

### FFU and immunofluorescence microscopy

Fluorescent focus unit (FFU) assays were used to determine the concentration of infectious virus in BKPyV purification or experimental samples. RPTE cells were infected with sample dilutions, fixed at 48 hpi and immunostained for VP1 expression as described in (Evans et al. 2015). Additionally, for **Figure 5C** infectious BKPyV levels of uninhibited conditions were arbitrarily set to 1 and inhibited conditions corrected to this control, 7 independent experiments. A one sample t-test was conducted to give *p* values, standard deviation shown with error bars.

For immunofluorescence analysis, RPTE cells were fixed in 3% formaldehyde. Fixed cells were then permeabilised and quenched (50mM NH_4_Cl and 0.1% Triton X-100 in PBS), blocked in PGAT (0.2% gelatin, 0.01% Triton X-100, 0.02% NaN_3_ in PBS) and stained with primary antibodies. Secondary antibodies used for immunofluorescence were Alexa Fluor 568 donkey anti-mouse or goat anti-IgG1 mouse and Alexa Fluor 488 donkey anti-rabbit or goat anti-IgG2a mouse. Coverslips were mounted using SlowFade Gold with DAPI (Invitrogen). Samples were imaged using a 63x oil immersion lens on an Olympus IX81 wide-field fluorescent microscope.

### Western blot

RPTE cells were lysed by suspending in mRIPA (50mM Tris pH 7.5, 150mM NaCl, 1% Sodium Deoxycholate and 1% Triton X-100) supplemented with Complete Protease Inhibitors without EDTA (Roche). Cellular debris were removed by centrifugation at 17,000g. HU cells were lysed by suspending in HU cell Lysis Buffer (20mM HEPES pH 7.6, 250mM Sucrose, 2mM DTT, 2mM EDTA Na_2_ and 2mM EGTA) supplemented with Complete Protease Inhibitors without EDTA, followed by sonication at 25W for 30 sec. Proteins were separated by SDS-PAGE electrophoresis and transferred to nitrocellulose membranes before blocking in 5% skimmed milk powder in PBS.

Following primary antibody binding, LI-COR IRDye680-(anti-mouse, anti-rabbit or anti-rat) or IRDye800-conjugated (anti-mouse or anti-rabbit) secondary antibodies were used. Membranes were then imaged on a LI-COR Odyssey Infrared Imaging system.

### Real-time PCR (qPCR)

RPTE cell pellets were lysed in 200µL NDA Lysis Buffer (4M Guanidine Thiocyanate, 25mM Tris and 134mM β-mercaptoethanol) and incubated at 56°C for 10 min, after which an equal volume of 100% ethanol was added. DNA was then bound to silica columns by centrifuging at 16,000× g for one min. Columns were washed with Buffer 1 (1M Guanidine Thiocyanate, 25mM Tris pH7 in 10% ethanol), and centrifuged, followed by a final wash in Buffer 2 (25mM Tris pH7 in 70% ethanol). DNA was eluted with nuclease free water by centrifugation at 16,000× g. Primers and probe for BKPyV genome were designed as described in (Evans et al. 2015). Human TNFα primers and probe were designed and obtained through TIB MOLBIOL (forward primer: AGGAACAGCACAGGCCTTAGTG; reverse primer: AAGACCCCTCCCAGATAGATGG; Taqman probe: CCAGGATGTGGAGAGTGAACCGACATG). 300nM of each primer and 50nM of Taqman probe were used in each qPCR reaction, run on a Rotor-Gene (RG-3000, Corbett Research) and subsequently analysed on Rotor-Gene software. BKPyV genome levels were corrected to the TNFα control for each sample, and uninhibited samples arbitrarily set to 1, 6 independent experiments. A one sample t-test was conducted to give *p* values, standard deviation shown with error bars.

### Flow cytometry

Cellular DNA content was used as an indicator of cell cycle status. Cells were fixed in 70% ethanol for 30 mins, DNA was stained by resuspending each PBS washed cell pellet in 0.2mg RNAse A and 50µg propidium iodide in 1mL PBS and incubated at 37°C for 1 hr. Cells were then centrifuged at 6,000 g, supernatant removed and resuspended in 500µL PBS. Cells were analysed by flow cytometry using BD FACSCantoII with BD FACSDiva Software (BD Biosciences) and further analysed using FlowJo v10.4.2 cell cycle analysis function. A minimum of 10,000 cells were collected for each sample, 3 independent experiments. Standard deviation error was calculated for each cell cycle status sample.

### Whole cell lysate protein digestion

Cells were washed twice with PBS, and 250 µλ lysis buffer added (6M Guanidine/50 mM HEPES pH 8.5). Cell lifters (Corning) were used to scrape cells in lysis buffer, which was removed to an eppendorf tube, vortexed extensively then sonicated. Cell debris was removed by centrifuging at 21,000 g for 10 min twice. Dithiothreitol (DTT) was added to a final concentration of 5 mM and samples were incubated for 20 mins. Cysteines were alkylated with 14 mM iodoacetamide and incubated 20 min at room temperature in the dark. Excess iodoacetamide was quenched with DTT for 15 mins. Samples were diluted with 200 mM HEPES pH 8.5 to 1.5 M Guanidine followed by digestion at room temperature for 3 h with LysC protease at a 1:100 protease-to-protein ratio. Samples were further diluted with 200 mM HEPES pH 8.5 to 0.5 M Guanidine. Trypsin was then added at a 1:100 protease-to-protein ratio followed by overnight incubation at 37°C. The reaction was quenched with 5% formic acid, then centrifuged at 21,000 g for 10 min to remove undigested protein. Peptides were subjected to C18 solid-phase extraction (SPE, Sep-Pak, Waters) and vacuum-centrifuged to near-dryness.

### Peptide labelling with tandem mass tags

In preparation for TMT labelling, desalted peptides were dissolved in 200 mM HEPES pH 8.5. Peptide concentration was measured by microBCA (Pierce), and 25 µg labelled with TMT reagent. TMT reagents (0.8 mg) were dissolved in 43 µl anhydrous acetonitrile and 3 µl added to peptide at a final acetonitrile concentration of 30% (v/v). Samples were labelled as follows. Experiment 1 (9-plex); 126 – mock infection 12 hpi, 127N – mock infection 24 hpi, 127C – mock infection 48 hpi, 128N – BKPyV infection 12 hpi, 128C – BKPyV infection 24 hpi, 129N – BKPyV irradiated – 48 hpi, 129C – BKPyV irradiated 24 hpi, 130N BKPyV infection 48 hpi. Experiment 2 (10-plex); 126 – HU cells mock infection 24 hpi, 127N – HU cells mock infection 72 hpi, 127C – HU cells BKPyV infection 24 hpi, 128N – HU cells BKPyV infection 48 hpi, 128C – HU cells BKPyV infection 72 hpi, 129N – RPTE cells mock infection 24 hpi, 129C – RPTE cells mock infection 72 hpi, 130N – RPTE cells BKPyV infection 24 hpi, 130C – RPTE cells BKPyV infection 48 hpi, 131N – RPTE cells BKPyV infection 72 hpi. Following incubation at room temperature for 1 h, the reaction was quenched with hydroxylamine to a final concentration of 0.3% (v/v). TMT-labelled samples were combined at a 1:1:1:1:1:1:1:1:1 ratio (experiment 1) and 1:1:1:1:1:1:1:1:1:1 ratio (experiment 2). The sample was vacuum-centrifuged to near dryness and subjected to C18 SPE (Sep-Pak, Waters). An unfractionated singleshot was initially analysed to ensure similar peptide loading across each TMT channel, to avoid the need for excessive electronic normalization. Quantities of each TMT labelled sample were adjusted prior to high pH reversed-phase (HpRP) so that normalisation factors were >0.67 and <1.5. Normalisation is discussed in ‘Data Analysis’, and fractionation is discussed below.

### Offline HpRP fractionation

TMT-labelled tryptic peptides were subjected to HpRP fractionation using an Ultimate 3000 RSLC UHPLC system (Thermo Fisher Scientific) equipped with a 2.1 mm internal diameter (ID) × 25 cm long, 1.7 µm particle Kinetix Evo C18 column (Phenomenex). Mobile phase consisted of A: 3% acetonitrile (MeCN), B: MeCN and C: 200 mM ammonium formate pH 10. Isocratic conditions were 90% A / 10% C, and C was maintained at 10% throughout the gradient elution. Separations were conducted at 45°C. Samples were loaded at 200 µl/minute for 5 minutes. The flow rate was then increased to 400 µl/minute over 5 minutes, after which the gradient elution proceed as follows: 0-19% B over 10 minutes, 19-34% B over 14.25 minutes, 34-50% B over 8.75 minutes, followed by a 10 minutes wash at 90% B. UV absorbance was monitored at 280 nm and 15 s fractions were collected into 96 well microplates using the integrated fraction collector. Fractions were recombined orthogonally in a checkerboard fashion, combining alternate wells from each column of the plate into a single fraction, and commencing combination of adjacent fractions in alternating rows. Wells prior to the start or after the stop of elution of peptide-rich fractions, as identified from the UV trace, were excluded. This yielded two sets of 12 combined fractions, A and B, which were dried in a vacuum centrifuge and resuspended in 10 µl MS solvent (4% MeCN / 5% formic acid) prior to LC-MS3. 11 set ‘A’ fractions were used for experiment 1 and 10 set ‘A’ fractions were used for experiment 2.

### LC-MS3

Mass spectrometry data was acquired using an Orbitrap Lumos (Thermo Fisher Scientific, San Jose, CA). An Ultimate 3000 RSLC nano UHPLC equipped with a 300 µm ID × 5 mm Acclaim PepMap µ-Precolumn (Thermo Fisher Scientific) and a 75 µm ID × 50 cm 2.1 µm particle Acclaim PepMap RSLC analytical column was used.

Loading solvent was 0.1% formic acid (FA), analytical solvent A: 0.1% FA and B: 80% MeCN + 0.1% FA. All separations were carried out at 55°C. Samples were loaded at 5 µL/minute for 5 minutes in loading solvent before beginning the analytical gradient. The following gradient was used: 3-7% B over 3 minutes, 7-37% B over 173 minutes, followed by a 4 minute wash at 95% B and equilibration at 3% B for 15 minutes. Each analysis used a MultiNotch MS3-based TMT method (McAlister et al. 2012, McAlister et al. 2014). The following settings were used Th, 120,000 Resolution, 2×10^5^ automatic gain control (AGC) target, 50 ms maximum injection time. MS2: Quadrupole isolation at an isolation width of m/z 0.7, CID fragmentation (normalised collision energy (NCE) 35) with ion trap scanning in turbo mode from m/z 120, 1.5×10^4^ AGC target, 120 ms maximum injection time. MS3: In Synchronous Precursor Selection mode the top 6 MS2 ions were selected for HCD fragmentation (NCE 65) and scanned in the Orbitrap at 60,000 resolution with an AGC target of 1×10^5^ and a maximum accumulation time of 150 ms. Ions were not accumulated for all parallelisable time. The entire MS/MS/MS cycle had a target time of 3 s. Dynamic exclusion was set to +/-10 ppm for 70 s. MS2 fragmentation was trigged on precursors 5×10^3^ counts and above.

### Data analysis

In the following description, we list the first report in the literature for each relevant algorithm. Mass spectra were processed using a Sequest-based software pipeline for quantitative proteomics, “MassPike”, through a collaborative arrangement with Professor Steve Gygi’s laboratory at Harvard Medical School. MS spectra were converted to mzXML using an extractor built upon Thermo Fisher’s RAW File Reader library (version 4.0.26). In this extractor, the standard mzxml format has been augmented with additional custom fields that are specific to ion trap and Orbitrap mass spectrometry and essential for TMT quantitation. These additional fields include ion injection times for each scan, Fourier Transform-derived baseline and noise values calculated for every Orbitrap scan, isolation widths for each scan type, scan event numbers, and elapsed scan times. This software is a component of the MassPike software platform and is licensed by Harvard Medical School.

A combined database was constructed from (a) the human Uniprot database (4th February 2014), (b) the BK polyomavirus database (6th October, 2014). The combined database was concatenated with a reverse database composed of all protein sequences in reversed order. Searches were performed using a 20 ppm precursor ion tolerance (Haas et al. 2006). Product ion tolerance was set to 0.03 Th. TMT tags on lysine residues and peptide N termini (229.162932 Da) and carbamidomethylation of cysteine residues (57.02146 Da) were set as static modifications, while oxidation of methionine residues (15.99492 Da) was set as a variable modification.

To control the fraction of erroneous protein identifications, a target-decoy strategy was employed (Elias and Gygi 2007, Elias and Gygi 2010). Peptide spectral matches (PSMs) were filtered to an initial peptide-level false discovery rate (FDR) of 1% with subsequent filtering to attain a final protein-level FDR of 1% (Kim et al. 2011, Wu et al. 2011). PSM filtering was performed using a linear discriminant analysis, as described previously (Huttlin et al. 2010). This distinguishes correct from incorrect peptide IDs in a manner analogous to the widely used Percolator algorithm (Kall et al. 2007), though employing a distinct machine learning algorithm. The following parameters were considered: XCorr, ΔCn, missed cleavages, peptide length, charge state, and precursor mass accuracy. Protein assembly was guided by principles of parsimony to produce the smallest set of proteins necessary to account for all observed peptides (Huttlin et al. 2010).

Proteins were quantified by summing TMT reporter ion counts across all matching peptide-spectral matches using “MassPike”, as described previously (McAlister et al. 2012, McAlister et al. 2014). Briefly, a 0.003 Th window around the theoretical m/z of each reporter ion (126, 127n, 127c, 128n, 128c, 129n, 129c, 130n, 130c, 131n, 131c) was scanned for ions, and the maximum intensity nearest to the theoretical m/z was used. The primary determinant of quantitation quality is the number of TMT reporter ions detected in each MS3 spectrum, which is directly proportional to the signal-to-noise (S:N) ratio observed for each ion (Makarov and Denisov 2009). Conservatively, every individual peptide used for quantitation was required to contribute sufficient TMT reporter ions (minimum of ∼1250 per spectrum) so that each on its own could be expected to provide a representative picture of relative protein abundance (McAlister et al. 2012). An isolation specificity filter was additionally employed to minimise peptide co-isolation (Ting et al. 2011). Peptide-spectral matches with poor quality MS3 spectra (more than 9 TMT channels missing and/or a combined S:N ratio of less than 250 across all TMT reporter ions) or no MS3 spectra at all were excluded from quantitation. Peptides meeting the stated criteria for reliable quantitation were then summed by parent protein, in effect weighting the contributions of individual peptides to the total protein signal based on their individual TMT reporter ion yields. Protein quantitation values were exported for further analysis in Excel.

For protein quantitation, reverse and contaminant proteins were removed, then each reporter ion channel was summed across all quantified proteins and normalised assuming equal protein loading across all channels. For further analysis and display in figures, fractional TMT signals were used (i.e. reporting the fraction of maximal signal observed for each protein in each TMT channel, rather than the absolute normalized signal intensity). This effectively corrected for differences in the numbers of peptides observed per protein. For all TMT experiments, normalised S:N values are presented in Table S1 (‘Data’ worksheet).

Hierarchical centroid clustering based on uncentered Pearson correlation, and k-means clustering were performed using Cluster 3.0 (Stanford University) and visualised using Java Treeview (http://jtreeview.sourceforge.net) unless otherwise noted.

## Author Contributions

L.G.C., R.A. and C.T.R.D. acquired and analysed the experimental data. L.G.C., C.T.R.D., P.J.L., M.P.W. and C.M.C. conceived and designed the experiments, interpreted the data and contributed to writing the manuscript. M.P.W. and C.M.C. supervised the project.

## Acknowledgments

This work was supported by an Isaac Newton Trust / Wellcome Trust ISSF award to C.M.C., a Biotechnology and Biological Sciences Research Council grant (BB/M021424/1) to C.M.C., a Medical Research Council funded PhD studentship to L.G.C. and a Wellcome Trust Senior Clinical Research Fellowship (108070/Z/15/Z) to M.P.W.

## Supplementary Data Table descriptions

**Table S1.** Interactive spreadsheet of all data in the manuscript. The “DATA” worksheet shows minimally annotated protein data, with only formatting and normalisation modifying the raw data. The “Plotter” worksheet enables generation of individual protein abundance changes for both viral and human proteins over time from experiments 1 and 2. The total number of quantified peptides and proteins from experiments 1 and 2 are shown in separate workbook sheets.

**Table S2.** A list of all quantified proteins from experiments 1 and 2 that match the UniProt keywords terms ‘Innate Immunity’ and ‘Antiviral Defense’ and their corresponding fold changes post-BKPyV infection.

**Table S3.** Proteins modulated during BK infection. All proteins downregulated or upregulated >2-fold at any time point during the course of infection (compared to the average of the mock samples). Results from experiments 1 and 2 are presented.

**Table S4.** DAVID functional enrichment analysis from proteins upregulated or downregulated >2-fold against a background of all proteins quantified. Only significant (Benjamini-Hochberg corrected) clusters from experiments 1 and 2 are shown. There were no significant clusters amongst proteins downregulated >2-fold in experiment 2.

